# TagSeqTools: a flexible and comprehensive analysis pipeline for NAD tagSeq data

**DOI:** 10.1101/2020.03.09.982934

**Authors:** Huan Zhong, Zongwei Cai, Zhu Yang, Yiji Xia

**Affiliations:** State Key Laboratory of Environmental and Biological Analysis, Department of Chemistry, Hong Kong Baptist University, Hong Kong, China; Department of Biology, Hong Kong Baptist University, Hong Kong, China

**Keywords:** Transcriptome, Oxford nanopore sequencing, NAD-tagged RNAs, NAD tagSeq, pipeline

## Abstract

NAD tagSeq has recently been developed for the identification and characterization of NAD^+^-capped RNAs (NAD-RNAs). This method adopts a strategy of chemo-enzymatic reactions to label the NAD-RNAs with a synthetic RNA tag before subjecting to the Oxford Nanopore direct RNA sequencing. A computational tool designed for analyzing the sequencing data of tagged RNA will facilitate the broader application of this method. Hence, we introduce TagSeqTools as a flexible, general pipeline for the identification and quantification of tagged RNAs (i.e., NAD^+^-capped RNAs) using long-read transcriptome sequencing data generated by NAD tagSeq method. TagSeqTools comprises two major modules, <u>TagSeek</u> for differentiating tagged and untagged reads, and <u>TagSeqQuant</u> for the quantitative and further characterization analysis of genes and isoforms. Besides, the pipeline also integrates some advanced functions to identify antisense or splicing, and supports the data reformation for visualization. Therefore, TagSeqTools provides a convenient and comprehensive workflow for researchers to analyze the data produced by the NAD tagSeq method or other tagging-based experiments using Oxford nanopore direct RNA sequencing. The pipeline is available at https://github.com/dorothyzh/TagSeqTools, under Apache License 2.0.

## INTRODUCTION

Several non-canonical initiating nucleotides (NCINs), such as NAD^+^, NADH, and 3’-dephospho-coenzyme A (dpCoA), have been reported to cap the 5’ end of RNA (Chen et al. 2009; Kowtoniuk et al. 2009). It has been discovered since the early 1970s that a messenger RNA (mRNA) of eukaryotes and certain eukaryotic viruses typically bears a 7-methylguanylate (m7G) cap at its 5’ end (Furuichi, Muthukrishnan, and Shatkin 1975; Wei and Moss 1975). Decades of study revealed that m7G cap plays an essential role in the coordination of various cellular processes throughout the whole life cycle of mRNA, including efficient pre-mRNA splicing in nuclei (Konarska, Padgett, and Sharp 1984; Inoue et al. 1989; Fresco and Buratowski 1996), nuclear export of mature mRNA (Visa et al. 1996), initiation of cellular mRNA translation (Kapp and Lorsch 2004), the formation of the pseudo-circular structure of translating mRNA (Thomas Preiss and Hentze 1998; T. Preiss, Muckenthaler, and Hentze 1998), preventing mRNA from exonuclease degradation (Gao et al. 2000; Evdokimova et al. 2001), and formation of a signature of self RNA (Schuberth-Wagner et al. 2015). The NCIN caps have been identified first in bacteria (Chen et al. 2009; Kowtoniuk et al. 2009; Cahová et al. 2015), and then in human cells (Jiao et al. 2017), yeast (Walters et al. 2017) and plants (Wang et al. 2019; Zhang et al. 2019). It was identified in the living systems that the 5’ ends of some RNAs were capped with NAD^+^, as well as other nucleoside-containing metabolites like NADH, FAD and dpCoA (Cahová et al. 2015; Bird et al. 2016). The *in vivo* occurrence of more NCIN caps, such as uridine-containing UDP-glucose, UDP-GlcNAc and MurNAc-pentapeptide, which have been observed *in vitro*, is still under confirmation (Julius and Yuzenkova 2019). Compare to the RNA with m7G cap, NAD-capped RNA (NAD-RNA) seems more universal in the living system, widely observed in both prokaryotes (Cahová et al. 2015; Frindert et al. 2018) and eukaryotes (Jiao et al. 2017; Walters et al. 2017; Wang et al. 2019; Zhang et al. 2019). However, little about the functions of the NAD cap, as well as other NCIN caps, have been characterized. Exploring the biological role of the NAD cap in the life cycle of RNA is of scientific interest, which highly depends on an efficient and accurate method to identify and quantify the NAD-RNA species.

Recently, we developed a new method, NAD tagSeq, for the analysis of NAD-RNA using the nanopore sequencing technology (Zhang et al. 2019). Nanopore sequencing technology is one of the third-generation sequencing (TGS) methods that Oxford Nanopore Technologies (ONT) developed. Like NAD captureSeq (Cahová et al. 2015), the provious procedure, NAD tagSeq uses the ADPRC-catalyzed enzymatic reaction and the CuAAC click chemistry reaction for tagging NAD-RNAs. However, instead of tagging with biotin, azide conjugating with a synthetic RNA tag is used in the click reaction. As a result, NAD-RNAs are tagged with the RNA tag through a 1,2,3-triazole linkage. The RNAs with and without the RNA tag are then directly sequenced using the Oxford nanopore single-molecule sequencing technology (Khoddami and Cairns 2013). Compared with the previous methods, such as NAD CaptureSeq (Cahová et al. 2015) and CapZyme Seq (Vvedenskaya et al. 2018), one of the vital improvements of NAD tagSeq is that this approach can directly sequence RNA with Oxford nanopore sequencer, circumventing RNA fragmentation, reverse transcription and PCR amplification (Khoddami and Cairns 2013). These advantages enable long-read single-molecule sequencing of NAD-RNA, resulting in more structural information about the entire RNA molecules. Enrichment of NAD-RNA is also not a necessary step in NAD tagSeq, providing the opportunity to compare the NAD-capped and uncapped RNA isoforms quantitatively.

To apply the NAD tagSeq method, we established a computer pipeline which we termed as “TagSeqTools”. TagSeqTools facilitated the discovery of more than 2000 NAD-RNA from Arabidopsis, which are abundant in the genes related to photosynthesis, protein synthesis, and responses to cytokinin and stresses (Zhang et al. 2019). TagSeqTools starts from the sequencing reads generated from regular basecallers for Oxford nanopore sequencing, such as Albacore and Guppy. The core functions of TagSeqTools are recognition of the tagged and untagged RNA reads, and quantification of the ratio of NAD cap, which are implemented by the tools TagSeek, and tagSeqQuant, respectively. For the function of mapping reads to a reference genome, the pipeline employs the well-applied tool Minmap2 to do it. We have also integrated additional functions for further analysis and visualization of the results. Applied on the published NAD tagSeq data of Arabidopsis Col-0 (Zhang et al. 2019), TagSeqTools provide a comprehensive and user-friendly analysis pipeline of NAD-RNA signatures of transcriptome landscape.

## RESULTS

### 1. Overview of the pipeline

The backbone of TagSeqTools (Figure 1) is straightforward. TagSeqTools first starts with nanopore basecallers to translate the raw current signals to RNA sequence reads; these reads are subsequentially classified as tagged and untagged RNA and mapped to the reference genome; the quantification of tagged and untagged RNA for each gene/isoform, and other further analyses of the results then come following. However, TagSeqTools is a flexible pipeline, which allows selection among the tools of redundant function, and application of various workflows. TagSeqTools employs the ONT’s basecallers, Albacore and Guppy (https://community.nanoporetech.com). Guppy is the default base-calling tool since a recent comparison showed that Guppy outperformed among the other tools with both excellent accuracy and high speed (Wick, Judd, and Holt 2019). However, Albacore is also included for an additional choice and comparison between different tools. All currently available software is unable to identify the non-canonical band between tagRNA and the tagged RNA sequences resulted from the NAD tagSeq experiment, we therefore develop a module, TagSeek, for the pipeline to distinguish the tagged and untagged RNAs based on the nanopore reads. The quantification of each gene or isoform can be subsequently given based on the mapped sequences with and without the tagRNA at their 5’ end. The pipeline also contains Minmap2 to map all identified tagged and untagged RNA sequences to reference genome, resulting in the unique alignments of full length reads. NAD tagSeq was designed for the quantitative measurement of the ratio of NAD cap since the experiment can be applied without purification of tagged RNA, and detect both tagged and untagged sequences simultaneously. The quantification tool, TagSeqQuant, is of importance for the pipeline to offering quantitative measurement of tagged RNA (i.e., NAD-RNA). TagSeqQuant also includes other functions to provide advanced analysis of the results, such as comparing isoforms derived from the same loci and detecting antisense transcripts or pre-mRNAs, the advantages of the long-reads sequencing offered by TGS platform.

**Figure 1.**
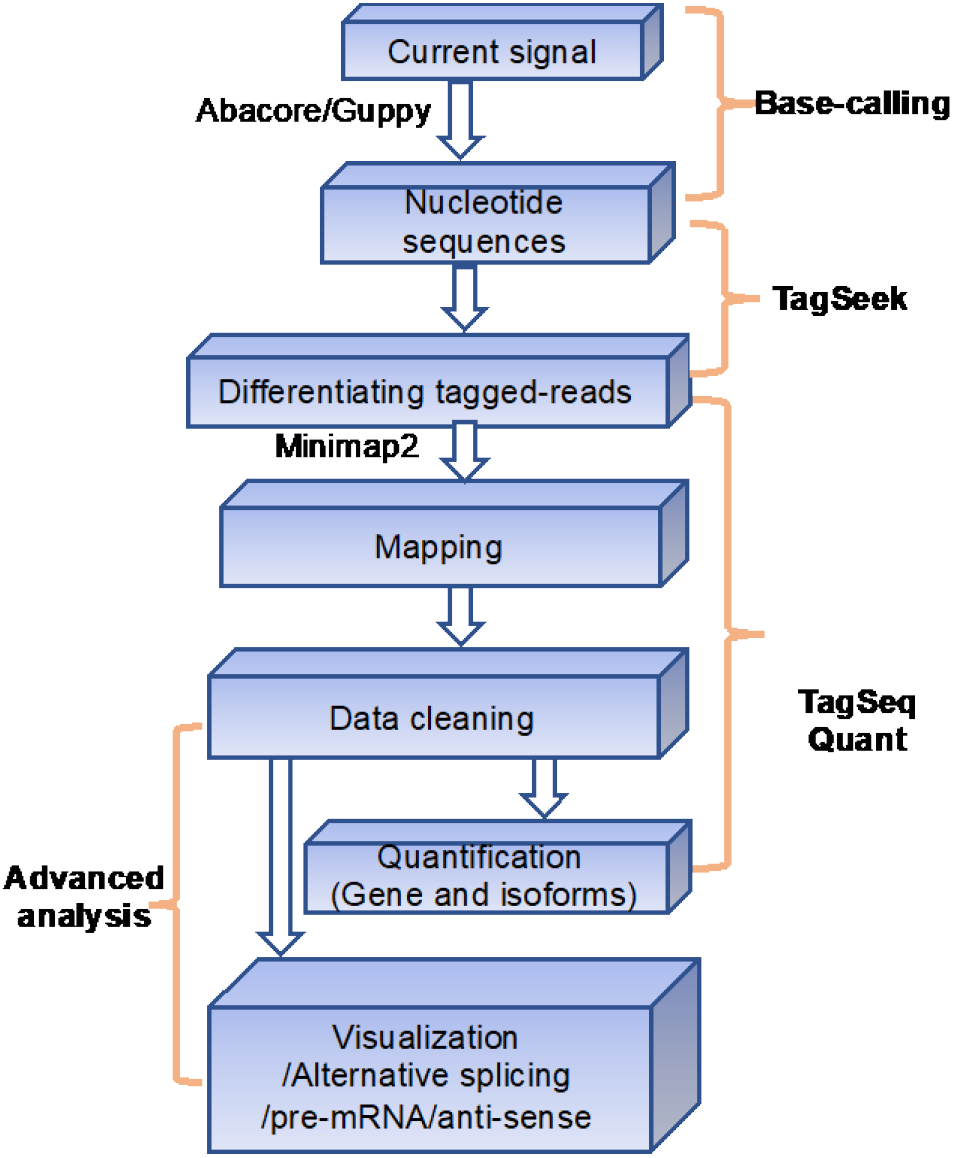
Schematic overview of the TagSeqTools pipeline. The pipeline was mainly divided into four parts: Base-calling (usually automatically finished when users use Nanopore sequencing); TagSeek module (mainly developed to differentiate tag and non-tag reads using the Nanopore fastq data); TagSeqQuant Module (integrating mapping, data cleaning, and quantification); and other independent codes for users’ interests in advanced analysis.

### 2. Classifying tagged and untagged RNAs using TagSeek

We first developed TagSeek to determine which reads are tagged and which is not since current basecallers are unable to call the linkage between synthetic tagRNA and NAD-RNAs directly. All reads were first grouped into two clusters based on the sequence matched to the tagRNA sequence in the 5’-end of each transcript. As illustrated in Figure 2A, these two groups of reads showed different patterns in coverage depth. The current version of the nanopore direct RNA sequencing method that is based on poly(A) selection and sequences RNA from 3’ end to 5’ end, resulting in 3’ bias. This phenomenon, which is typical for nanopore RNA sequencing, attributes to the trunked reads due to several possibilities, such as unexpected fragmentation during experiments or death of the nanopore before whole molecules pass (Khoddami and Cairns 2013; Keller et al. 2018).

**Figure 2.**
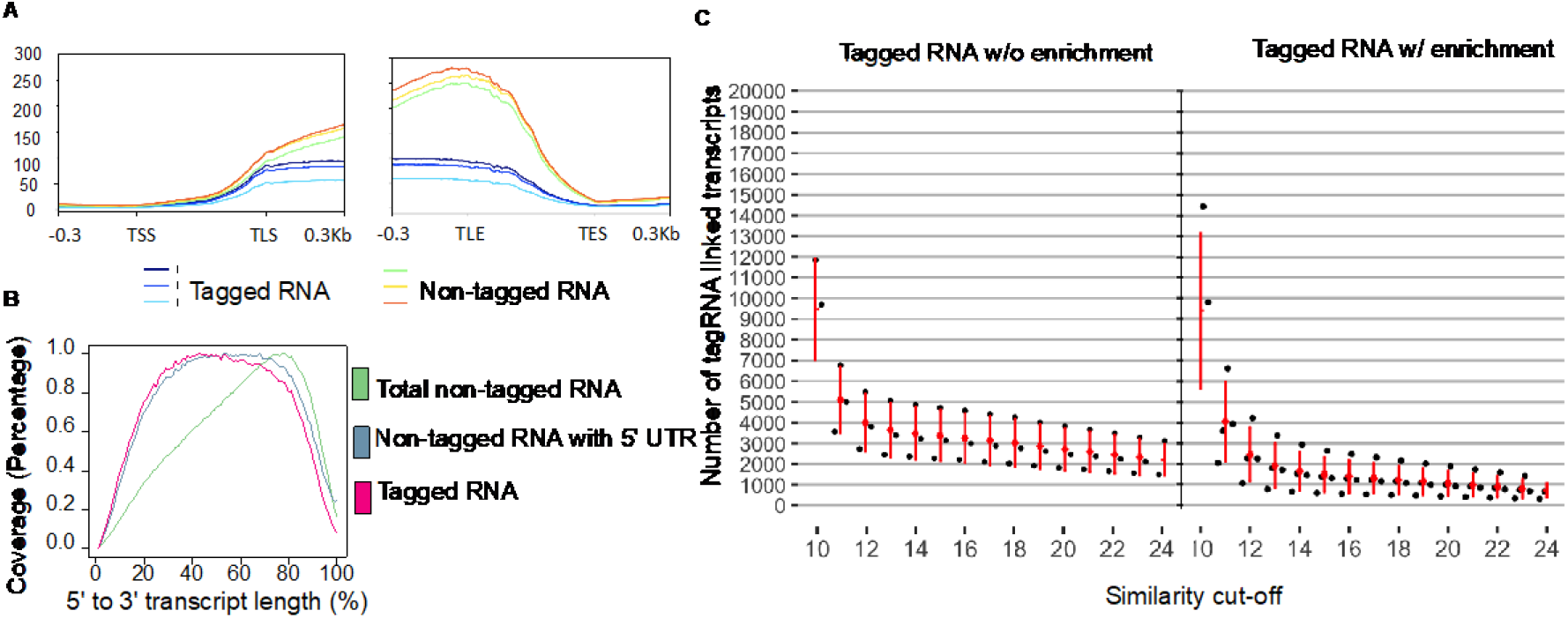
An evaluation of tagged RNA detection of TagSeqTools. A. Coverage depth over the RNA sequences with and without tagRNA at their 5’ end, including the transcription start site (TSS), transcription end site (TES), translation start site (TLS), and translation end site (TLE). B. The normalized mean sequencing coverages were calculated for the RNAs tagged in their 5’ UTRs (red), all non-tagged RNAs (green), and the non-tagged RNAs with their 5’ UTRs (blue). The length of each sequence was normalized according to the total length from the transcription start site (TSS) to the transcription end site (TES), and the coverage was normalized by dividing the maximum nucleotide read depth along with the sequences. C. Threshold selection for filtering of tagged RNAs. Left panel demonstrates the threshold selection for NAD tagSeq samples without tagged RNA enrichment, and right panel was for NAD tagSeq samples with tagged RNA enriched by hybridization to the DNA probe. The X-axis is similarity cut-off, namely the consecutively matched bases between tag sequence and first 40 bases of reads. The Y-axis is the number of tagged RNA transcripts passing specific similarity cut-off.

All trunked reads contribute only to the group without tag RNA, while the match of tag RNA in the 5’ end implicates the reads of full length. We, therefore, introduced the “symmetric” principle into TagSeqTools, requesting that untagged RNA reads should cover the same range at the 5’ ends as tagged RNA does. It leads to a similar coverage density of genes between tagged and untagged RNA reads (Figure 2B), avoiding the bias in determining both categories of RNAs.

TagSeek identifies the tagged RNA based on the sequence alignment between the tagRNA and sequences in the 5’-end of each RNA read. A 40 bases tagRNA was employed in the current version of NAD tagSeq. Tagseek, therefore, uses the first 40 bases of reads to map constitutively to the tagRNA sequence. It has been reported that nanopore sequencing has a dramatic loss of coverage in the ~10 nucleotides at the end of the sequences (here is the 5’ end of RNA since direct nanopore RNA sequencing starts from 3’ end) (Keller et al. 2018). Lower accuracy has also been observed in the ~10 nucleotides near the 1,2,3-triazole linkage due to the application of the basecallers that were designed for calling canonical nucleotides (Zhang et al. 2019). Thus, the maximum of matched sequences is 24 bases, much shorter than the tagRNA. Setting the various numbers of similarities to filter tagged RNA, we evaluated the results of different thresholds. For the cutoff less than 12, the average number of selected tagRNA-linked transcripts increased rapidly with the loosening of the cutoff, whereas the resulted dataset of tagged RNA changed a litter when the cutoff increased from 12 to 24 (Figure 2C). We, therefore, use 12 as the default threshold of the tagRNA match.

Taken together these criteria, TagSeek identified a read as tagged RNA if and only if it contained i) at least 12 bases at the 5’ end matching the tagRNA sequence, and ii) a main body of sequence mapped to a full-length reference sequence, i.e., from 5’ UTR to 3’ UTR.

### 3. Quantification of NAD-RNA and non-NAD-RNA using TagSeqQuant

NAD tagSeq uses ADPRC- samples as a negative control to reduce the false positive in tagged RNAs (Zhang et al. 2019). In addition to the arbitrary cutoff for TPM used in the previous work (Zhang et al. 2019), TagSeqTools also included Fisher’s exact test to estimate the significance of NAD-RNA levels in ADPRC+ samples compared to that in ADPRC- samples (Figure 3A). TagSeqQuant further calculated q-values using Benjamini–Hochberg adjustment based on the p-values calculated from Fisher’s exact test to correct the multiple testing errors. A cutoff of q-values < 0.1, for example, gave ~70% (3367 versus 2000) more NAD-RNA associated genes than the previous reports did (Figure 3B). Furthermore, the significance of these results was statistically sound.

**Figure 3.**
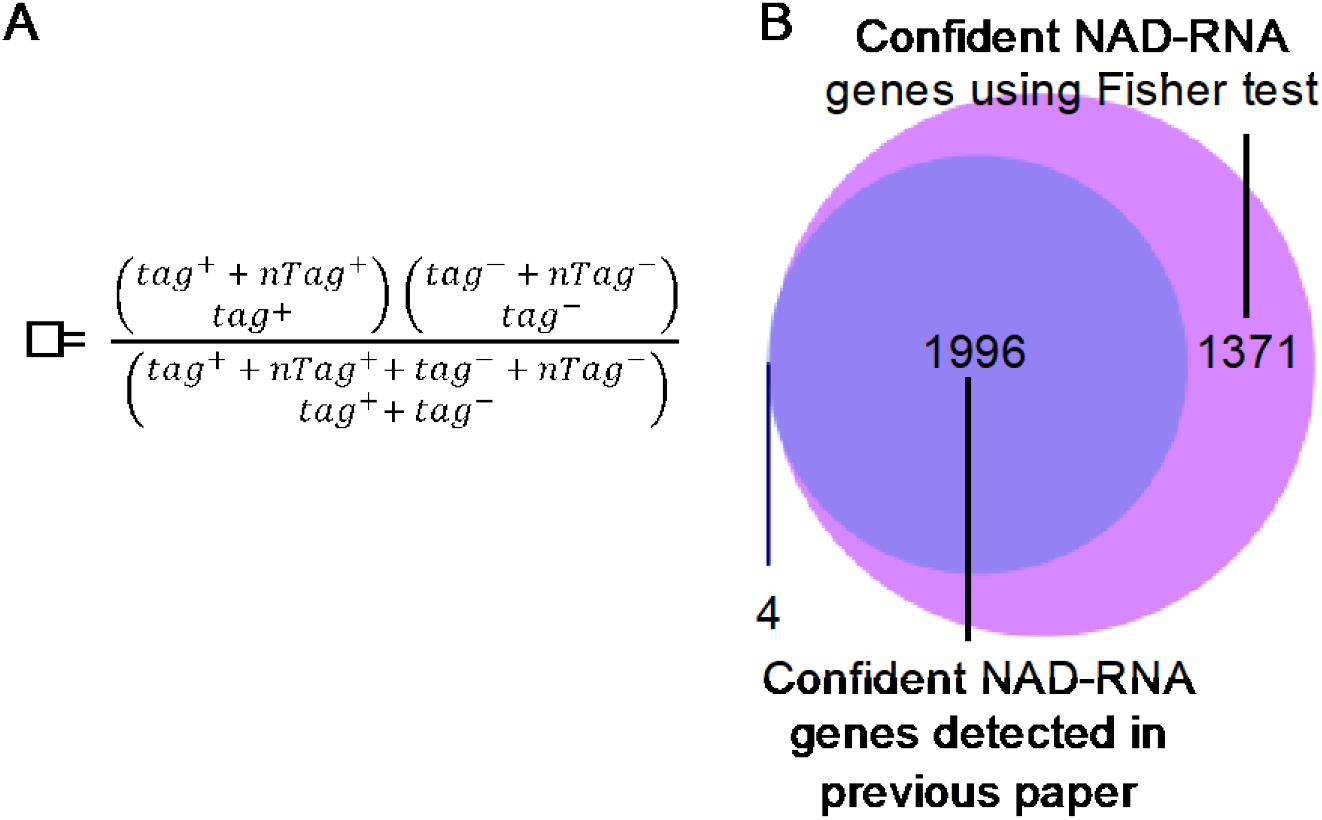
Quantification of NAD-RNA. A. The significance of NAD-RNA ratios was calculated using the Fisher exact test, based on the counts of tagged RNA in ADPRC+ (tag^+^) and ADPRC− (tag^−^) samples, as well as the counts of non-tagged RNA in the same parallel samples (nTag^+^ and nTag^−^, respectively). B. The comparison of the confident NAD-RNA genes found using the Benjamini–Hochberg adjusted Fisher test (q-value <0.1 in at least one replicate) to the published results.

**Figure 4.**
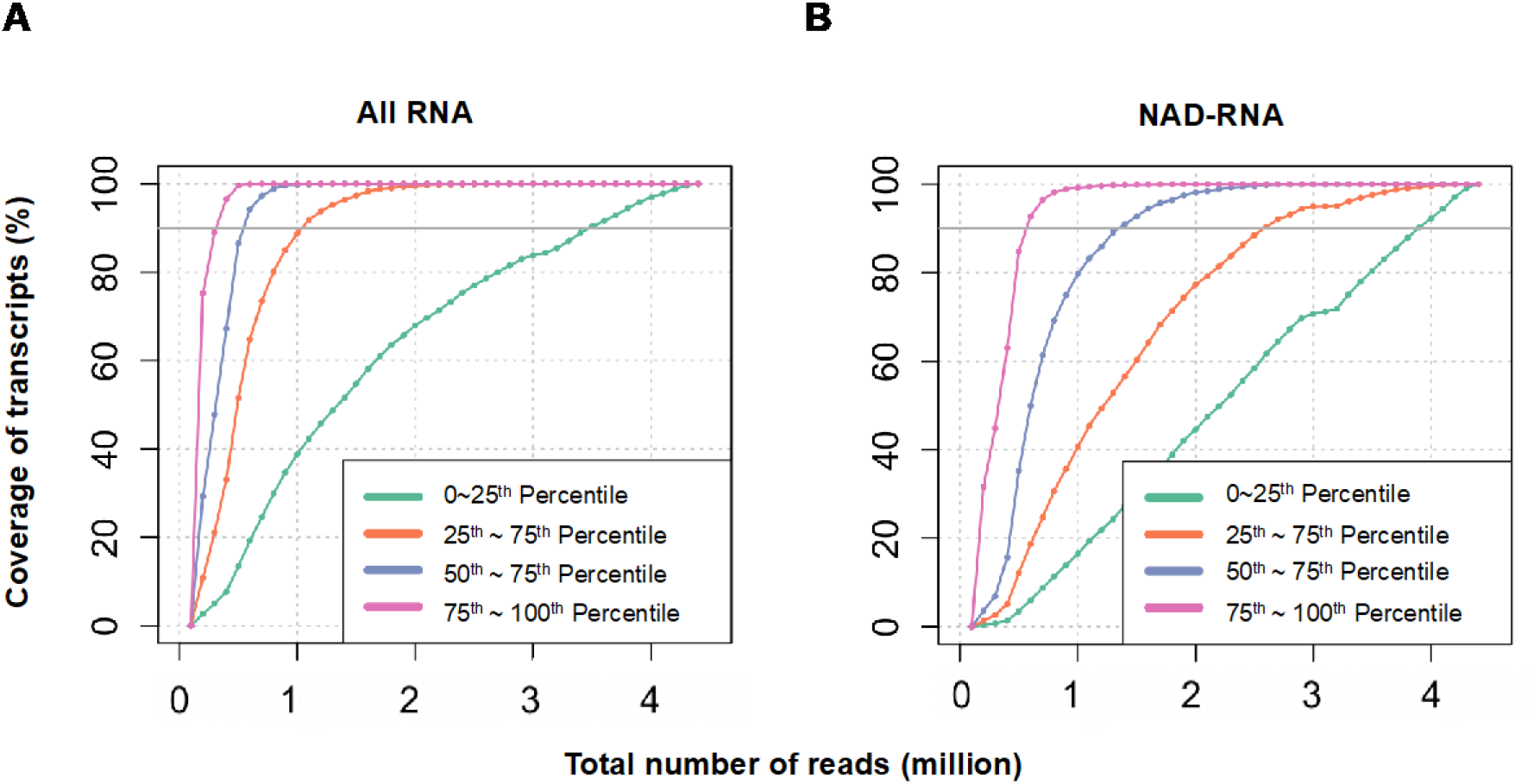
Gene expression levels at different sequencing depths. Quantification of transcripts levels from all genes (A) and NAD-RNA associated gene (B) at different sequencing depths. The statistics of the “final” TPM level were obtained at a depth of 4.25 million reads. Genes were sub-grouped based on their expression quantiles at the “Final” sequencing depth level. The 0~25^th^ percentile represents genes with the lowest expression levels, and the 75~100^th^ percentile represents genes with the highest expression levels. In each subset, the y-axis is the number of genes detected within the corresponding expression group for a certain number of reads, compared to the maximum number of genes detected in this group at the “Final” TPM level.

### 4 Transcript coverage with sequencing depth

We also constructed saturation curves to evaluate if the sequencing procedure reaches enough depth. We resampled the raw reads from the combination of three replicates and determined the number of genes detected in each pool. For all genes, 100% of the high abundance (in the top 25^th^ percentile) and 40% of the low abundance (in bottom 25^th^ percentile) genes were detected using 1 million reads (**Figure 3A**). For the detection of NAD-RNA associated genes, 100% of high abundance genes and only 20% of low abundance genes were detected using 1 million reads (**Figure 3B**). We show that 2.5 million reads are sufficient to detect 7362 genes with higher NAD-RNA levels (50^th^ ~100^th^ percentile; count > 5 and TPM > 1.41). The detection of genes (n = 3681) within the 25^th^~50^th^ percentile of expression level (count > 2, TPM > 0.56) almost approached saturation at roughly 4 million reads. Higher sequencing depth will be required in the future for the detection of genes with low abundance NAD-RNA transcripts. However, generally speaking, TagSeqTools can uncover signatures of NAD+ modified transcriptome with a relatively small number of sequencing reads.

### 5. Comparison of splicing patterns between NAD-RNA and non-NAD-RNA

The criteria used in TagSeek guarantees that TagSeqTools analysis results in full length reads of the transcripts, including the 3’ poly(A) tail and 5’ cap. This single-molecular RNA long-read sequencing approach allows the revealation of the association between NAD cap and the structural variants derived from a gene. Compared to traditional Illumina sequencing method, Nanopore sequencing has change to detect a whole transcript in one read. Thus isoforms can be much more easily recognized according to the difference of their structures (Supplementary Figure 1). From the overall comparison between the tagged and untagged RNAs, NAD-RNAs showed no significant difference in splicing manner from other RNAs at the total levels (Figures 5A - 5C). However, when we analyzed that at the individual level, there were 550 cases from Arabidopsis, of which different isoforms showed distinct ratios in NAD-capped transcripts. The filtering criteria are that i) variance of NAD-RNA ratio larger than 0.01, and ii) the counts of max NAD-RNA ratio of isoforms larger than 4. The top ten genes with varied NAD-RNA ratio in their isoforms were shown in Table 1. Most of these 550 genes (67%) and the top ten ones had two isoforms., However, some genes with more isoforms showed complicate patterns in the variance of NAD-RNA ratio among different types of transcripts (Figure 5D and 5E).

**Table 1.** The top ten genes with varied NAD-RNA ratio in their isoforms.

| Gene | Variance of NAD-RNA ratio in isoforms | Counts of max NAD-RNA ratio of isoforms | The max NAD-RNA ratio of isoforms | Counts of minimum NAD-RNA ratio of isoforms | The minimum NAD-RNA ratio of isoforms | Annotation |
| --- | --- | --- | --- | --- | --- | --- |
| GLP10 | 0.07<br>(2 isoforms) | 33 | 0.37 | 0 | 0.00 | Germin-like protein subfamily 2 member 4 |
| SCPL51 | 0.07<br>(2 isoforms) | 7 | 0.37 | 0 | 0.00 | Serine carboxypeptidase-like 51 |
| CYTB5-B | 0.06<br>(2 isoforms) | 196 | 0.35 | 0 | 0.00 | Cytochrome b5 isoform B |
| GCL1 | 0.06<br>(2 isoforms) | 8 | 0.35 | 0 | 0.00 | LanC-like protein GCL1 |
| GASA5 | 0.06<br>(2 isoforms) | 25 | 0.35 | 0 | 0.00 | Gibberellin-regulated protein 5 |
| SNRNP65 | 0.06<br>(2 isoforms) | 6 | 0.33 | 0 | 0.00 | U11/U12 small nuclear ribonucleoprotein 65 kDa protein |
| AT1G17640 | 0.06<br>(2 isoforms) | 5 | 0.33 | 0 | 0.00 | RNA-binding (RRM/RBD/RNP motifs) family protein |
| AT5G42680 | 0.05<br>(2 isoforms) | 15 | 0.33 | 0 | 0.00 | Protein of unknown function, DUF617 |
| AT1G06630 | 0.05<br>(2 isoforms) | 8 | 0.32 | 0 | 0.00 | F-box/LRR-repeat protein |
| AT1G18440 | 0.05<br>(2 isoforms) | 8 | 0.32 | 0 | 0.00 | Peptidyl-tRNA hydrolase, chloroplastic |

**Figure 5.**
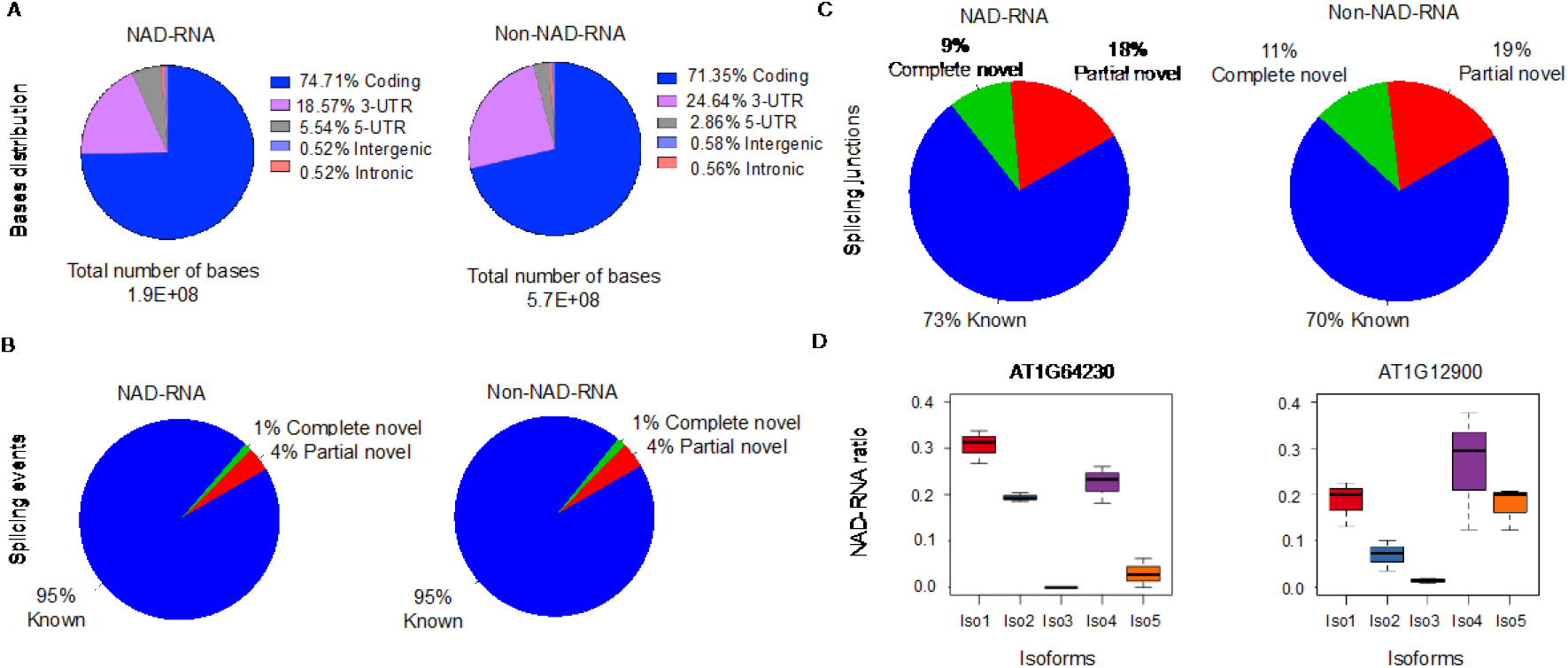
The NAD-capped isoforms resulted from alternative splicing. A. Distribution of bases mapped to different regions of the whole genome. B. Splicing junctions detected in NAD-RNA and non-NAD-RNA. C. Splicing events detected in NAD-RNA and non-NAD-RNA. D. Two example boxplots of the NAD-RNA ratio varied in different isoforms of the corresponding genes.

### 6. Polyadenylation in transcription starting sites of NAD-RNA

We estimated the distribution of ATCG nucleotides in the beginning of NAD-RNA reads. We found about 20 initial bases were consistent in nucleotide frequency, owing to the artificially added tag-sequence shall be somewhat similar among all the NAD-RNA reads, despite some mismatches (Figure 6A). Moreover, there was a sharp peak around 20^th^ nucleotides, followed by a relatively high A proportion in the following positions.

**Figure 6.**
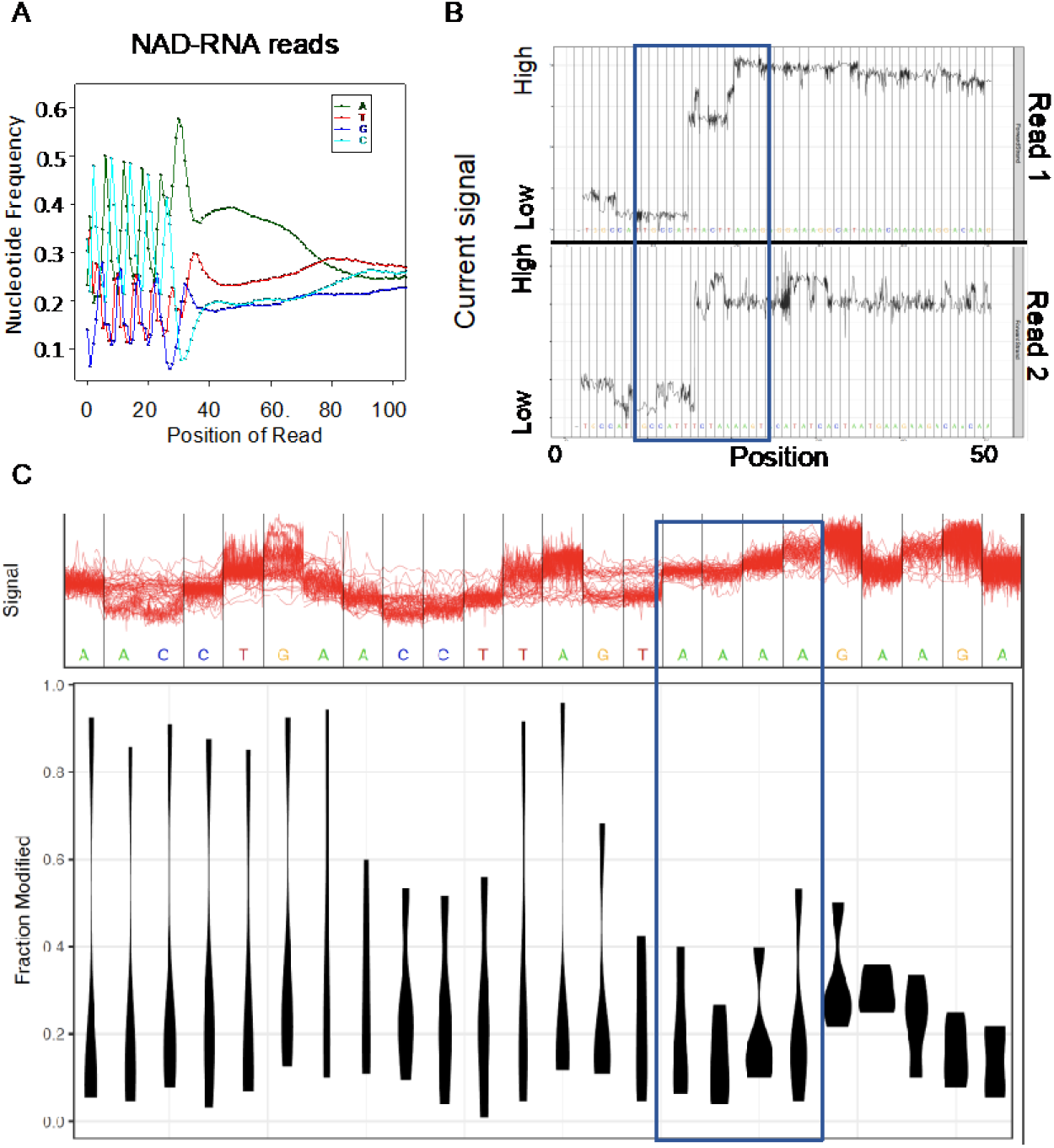
The linkage between tagRNA sequence and native RNA affects current signals. A. The first 100 bp of the tagged reads were used to calculate the nucleotide frequency. B. Examples of current signal plots in the first 50 bases of reads. Two reads were selected to represent the aberrant current signal fluctuation around the linkage between synthetic tag and NAD-RNA. The position highlighted in the blue box may refer to the linkage position. C. The detection of modified nucleotides. The raw genome-anchored nanopore signal in the top panel along with corresponding nucleotides and distribution of negative log p-values from the Fisher test of each base is shown in the below.

Furthermore, we also found the 1,2,3-triazole linkage may strongly disturb the current signals, with also a sharp fluctuation of the current signal around 20 bp upstream of the reads, and multiple consecutive A nucleotides observed in the disturbed region (Figure 6B). Besides, Tombo (Stoiber et al. 2017) was applied to estimate the modified bases of the reads using a machine learning method. Bases around the poly-A region were shown to have relatively higher p-values than other regions (Figure 6C), indicating they are more likely to be modified.

In order to assess whether the chemical linkage will disturb the current signal, which may be prone to be transformed into A nucleotides, or the modification of bases is more likely to happen in A-enriched regions, we perform the motif analysis on the 5’ UTR regions of genes. Frequency of AT nucleotides was quite higher in the annotated 5’ UTR regions, and the proportion of A and T were similar in the first 100 bp of 5’ UTR (Supplementary Figure 2). We selected the longest 5’ UTR of NAD-RNA read for each gene and referred the initiating site as the putative transcription start site (TSS). By extracted the ± 50 bp up- and downstream of the putative TSS from the annotated cDNA reference, we found a large portion of A-rich sites and TATA box located at the beginning of NAD-RNA transcripts. The results here implied that the actual NAD-RNA transcriptions were alternative and preferred to start from “A” nucleotides enriched regions.

## DISCUSSION

The NAD-tagSeq has distinct advantages for identification of NAD-capped RNAs (Zhang et al. 2019), and may indeed be considered the method of choice in the field for genome-wide NAD-capped RNA research. TagSeqTools was designed for processing the data generated from this protocol and facilitate the performance of the methods in broad applications. By combining the currently available tools such as basecallers for nanopore sequencing and the specific functional modules, TagSeqTools achieved the goal without developing a new basecaller specific to the 1,2,3-triazole linked RNA sequence. However, a shortage of the method is in determination of exact 5’ ends of NAD-RNAs. Such a basecaller that can recognize the sequence surrounding the 1,2,3-triazole linkage may be a useful extension of the pipeline. However, the basecalling from raw nanopore data is virtually machine-learning-based, which requires a large amount of data (Wick, Judd, and Holt 2019). Thus, it is hard to see such a tagged RNA-specific basecaller in the near future. The strategy used in TagSeqTools is arguable the best choice currently. Moreover, once another reaction to selectively attach the tagRNA to an NCIN was found, the NAD tagSeq method can be adapted easily to the NCIN cap with slight modifications. This new reaction results possibly in a different linkage between tagRNA and the target RNAs. TagSeqTools can be applied directly to data of the new experimental approach without modification. The specific basecaller, however, needs data accumulation and training from the very beginning.

In addition to the data procession for NAD tagSeq, TagSeqTools has integrated various of tools for further analyses, mainly based on the advantages of the NAD tagSeq method. One advantage of NAD tagSeq is direct sequencing of transcripts, skipping purification and amplification steps, and allowing quantification of relative abundance of of both NAD-capped and uncapped RNAs in a sample. TagSeqTools utilizes this information to calculate not only the ratio of NAD-RNA in each transcript (Figures 5D and 5E), which is biologically meaningful but also the p-values (Figure 3A), estimating the statistical significance of the quantification results. Another advantage of NAD tagSeq is that the nanopore sequencing technology, which allows RNA reads covering the full length of the molecules from 3’ end to 5’ end. This offers one an opportunity to study the association between NAD cap and structural variants derived from the same locus. The analysis of Arabidopsis data, for example, has revealed the cases of which the isoforms of the same genes have various ratios of NAD-capped transcripts (Tabel 1; Figures 5D and 5E). This is likely to suggest that the NAD cap may have a role in regulating alternative splicing of RNA, which is worth further investigation.

Besides the analysis at the transcriptomics level, studies from the sequence near TSS are another promising way to get insights into the regulations and/or functions of the NAD cap. Although failed at a few bases, TagSeqTools provide the information about the rough position of the NAD-capped nucleotides. The discovery of several motifs from the upstream region of NAD-capped nucleotides is very likely to have some functional roles. Although these sequences may be responsible for some regulatory regions such as promoters or silencers (Figure 7), the possibility that the formation of NAD-capped is also possibly regulated by one or more of these regions is still open. Further studies, such as comparison that to the same region of genes that produce more non-NAD-RNAs, or functional studies with genetic approaches, are necessary to answer this question. TagSeqTools, at the current stage, helps discover the candidate for the following considerations.

**Figure 7.**
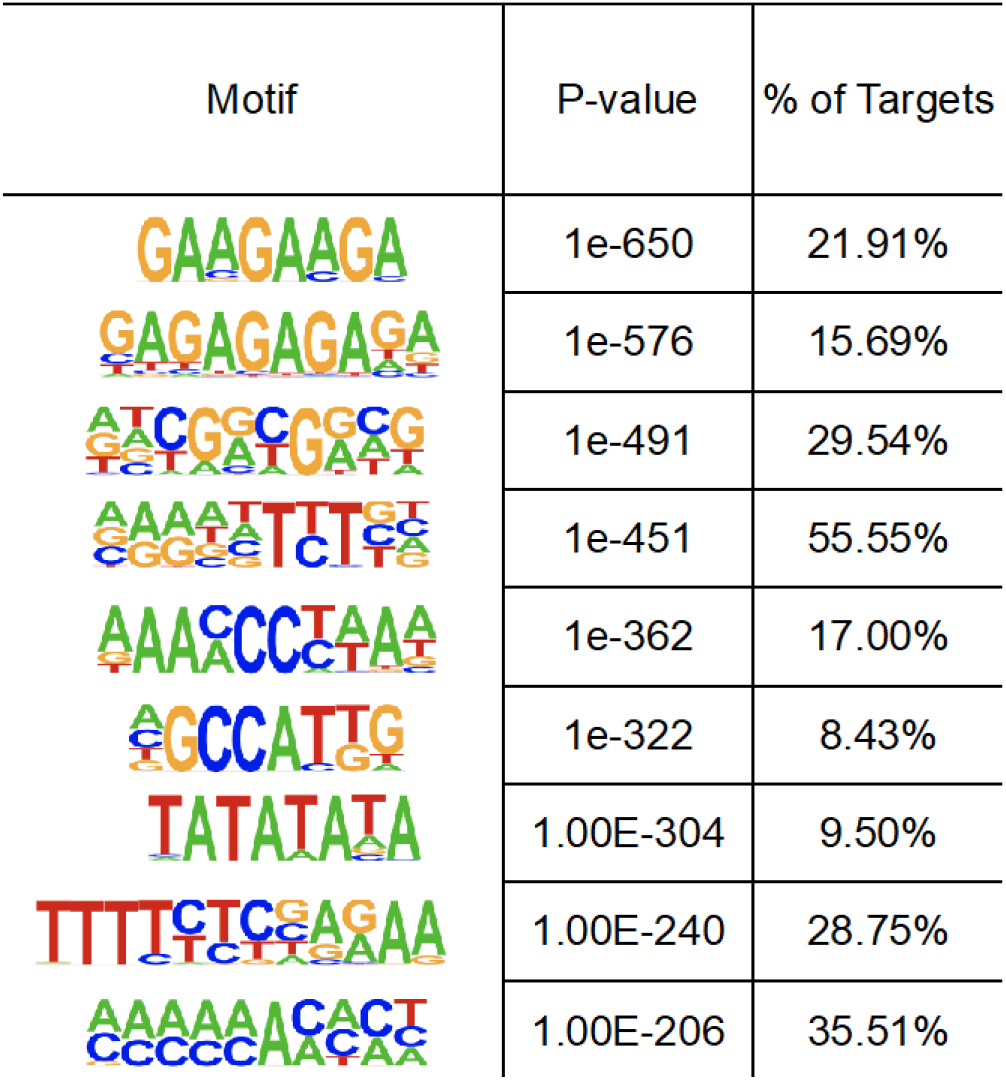
Potential motifs in NAD-RNA transcripts. Motifs were detected from the ± 50 bp surrounding the transcription starting site of NAD-RNAs along the annotated cDNA sequence using HOMER. The starting sites of NAD-RNAs were estimated based on the longest NAD-RNA transcripts of the corresponding gene.

## METHOD

### Data Availability and description

Current signal data (fast5) will be deposited in the GEO database. For the published dataset, RNA samples from 12-year-old Col-0 seedling of Arabidopsis were used. All samples were prepared using the NAD tagSeq method and sequenced by Nanopore direct RNA sequencing method. Raw fastq data were retrieved from (Khoddami and Cairns 2013) (accession no. GSE127755). tagRNA sequences were 40 nucleotides with “CCUGAA” repeats: 5’-CCUGAACCUGAACCUGAACCUGAACCUGAACCUGAACCUG/3AzideN/-3’.

Samples from the two experiments were included:

1. Tag RNA w/ enrichment: RNAs with native poly(A) tails were enriched from total RNA Col-0 and subjected to tagging with tagRNAs using ADPRC. After removal of free tagRNAs and further enrichment of poly(A)-containing RNAs using oligo (dT) beads, tagged RNAs were then enriched by hybridization to the DNA probe. A parallel experiment without ADPRC served as a negative control. Three biological replicates were included for both the ADPRC+ and ADPRC− samples.
2. Tag RNA w/o enrichment: An experiment similar to above but skipped the hybridization-based enrichment step for tagRNA-linked RNAs so that all transcripts from the same genes could be counted. Four samples included three biological replicates of ADPRC+ and one ADPRC−.

### TagSeek module

TagSeek module aimed to differentiate tagged or non-tagged reads from raw fastq files according to users’ preference on how many bases should be consecutively matched with tagRNA sequences (−t), which can be referred as tagged-reads. As tagRNA sequences in demo were repeats of 6 nucleotides “CCUGAA”, when similarity cut-off (−s) was set to be 12, there will be six kinds of 12-mer patterns, and TagSeek module will automatically select the reads which were including at least one pattern, write to additional two fastq files (*tag.fastq and *nontag.fastq). The number of reads matched to each pattern and statistics of results will also be shown in the final.

### TagSeqQuant module

The basic module integrates quality control, mapping, data cleaning and quantification steps.

#### 1) Quality control

FastQC (Wingett and Andrews 2018) was used to evaluate the quality of reads, including quality scores across all bases, GC content per base, sequence duplication levels and so on. Other quality control software can also be employed to implement the function.

#### 2) Mapping

Tagged-reads and nontagged-reads produced in the TagSeek were aligned to the genome and transcriptome fasta files separately using Minimap2 (v2.12) (Li 2018). We used genome mapping results for visualization and transcriptome mapping results for quantification.

#### 3) Data cleaning and quantification

A Rscript named TagSeqQuant.r was integrated into the TagSeqQuant module, which can perform data cleaning and quantify the number of tagged reads and non-tagged reads.

Reads in tagged or nontagged mapping results with multiple alignments were re-adjusted separately in this module, and the longest aligned gene/isoform in was kept for each read. Besides, tagged-reads which were aligned from the beginning of the native RNA were discarded for accuracy. 5’ ends of a large number of RNAs may be degraded through Nanopore direct RNA-sequencing. Therefore, the TagSeqQuant module can also allow users to keep full-length RNA transcripts covering 5’ to 3’ UTR.

The number of genes and isoforms were quantified for tagged reads and nontagged- eads using the same module and the statistics and counts tables will be produced for further analysis. In this pipeline, we added up the number of tagged-reads and nontagged-reads to define the total number of reads for each gene. Moreover, as reads can be mapped to the transcriptome in sense or antisense direction, the antisense aligned reads should be separately quantified for the downstream analysis.

### Advanced analysis

After all aforementioned steps, in this demo, we considered transcripts with tagRNA sequences were NAD-RNAs and transcripts without tagRNA sequences were non-NAD-RNAs. Raw counts aligned to each gene were normalized in TPM (Transcripts Per Million Reads) using the total number of unique mapped reads of the corresponding samples. Fisher’s Exact Test was performed in individual genes based on the number of NAD-RNA and non-NAD-RNA counts in ADPRC+ and ADPRC− samples, and FDR were obtained using p-values from the fisher tests. NAD-RNA ratios were calculated using NAD-RNA counts versus total aligned counts for each gene, in Tag RNA w/o enrichment samples, the NAD-RNA ratios can be regarded as NAD-RNA enriched ratio. These steps were all wrapped into an R script (advanced.r)

### Analysis of NAD-RNA transcription starting sites

Tombo 1.3 (Stoiber et al. 2017) was used for the current signal exploration and visualization, the fast5 files were re-squiggled first, before conducting further statistical analysis. Then Homer (Heinz et al. 2010) was applied for motif searching, and the top significant motifs were shown.

## SUPPLEMENTARY INFORMATION

**Supplementary Figure 1.**
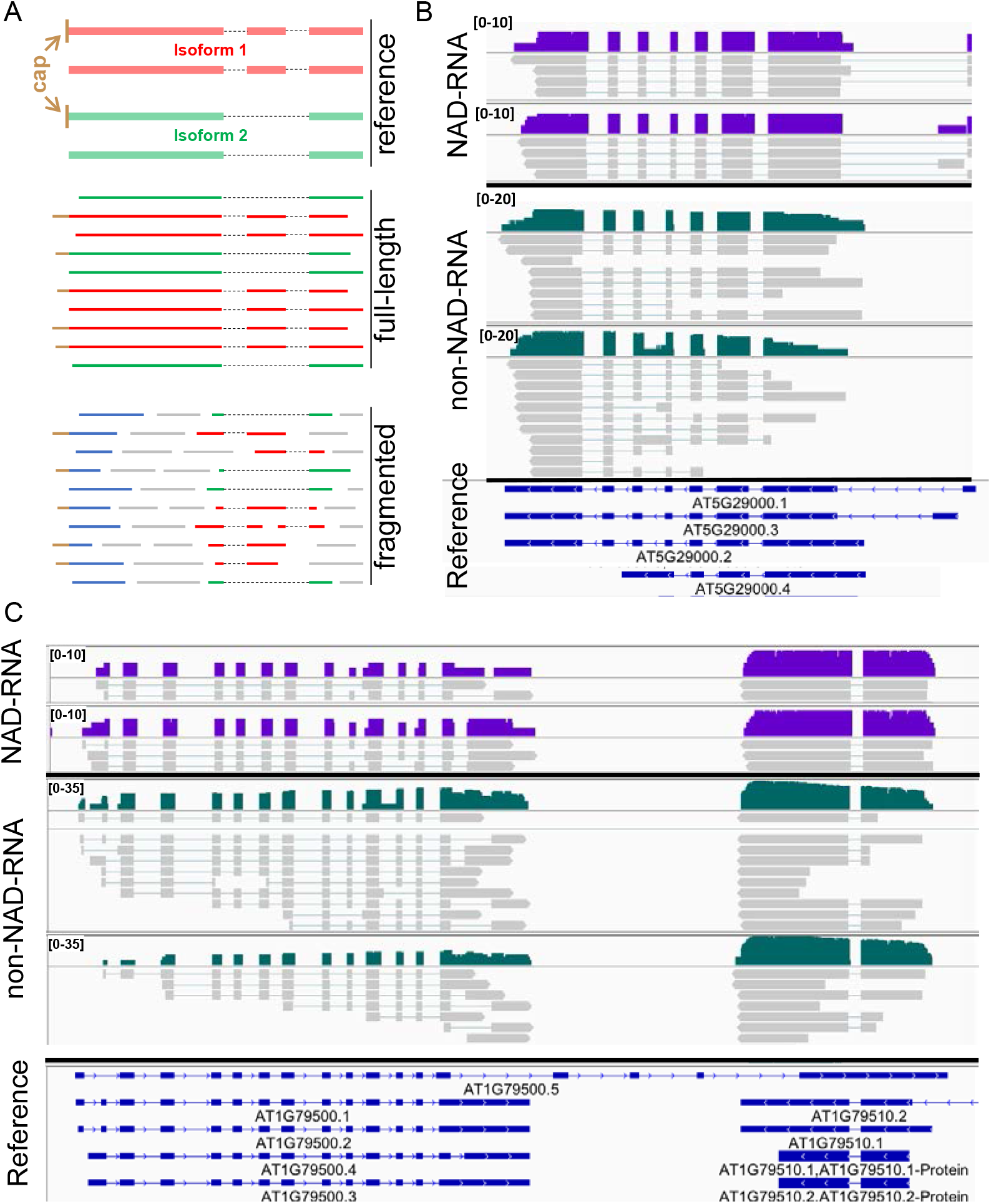
Isoform detection through long-read sequencing method. A. Schematic comparisons between Nanopore direct RNA sequencing and Illumina sequencing. B and C. Two examples of isoforms visualization plots. Read coverage (log scale in IGV) of two replicates for NAD-RNA (deep purple) and non-NAD-RNA (darkgreen) were shown. Individual reads were displayed in grey color.

**Supplementary Figure 2.**
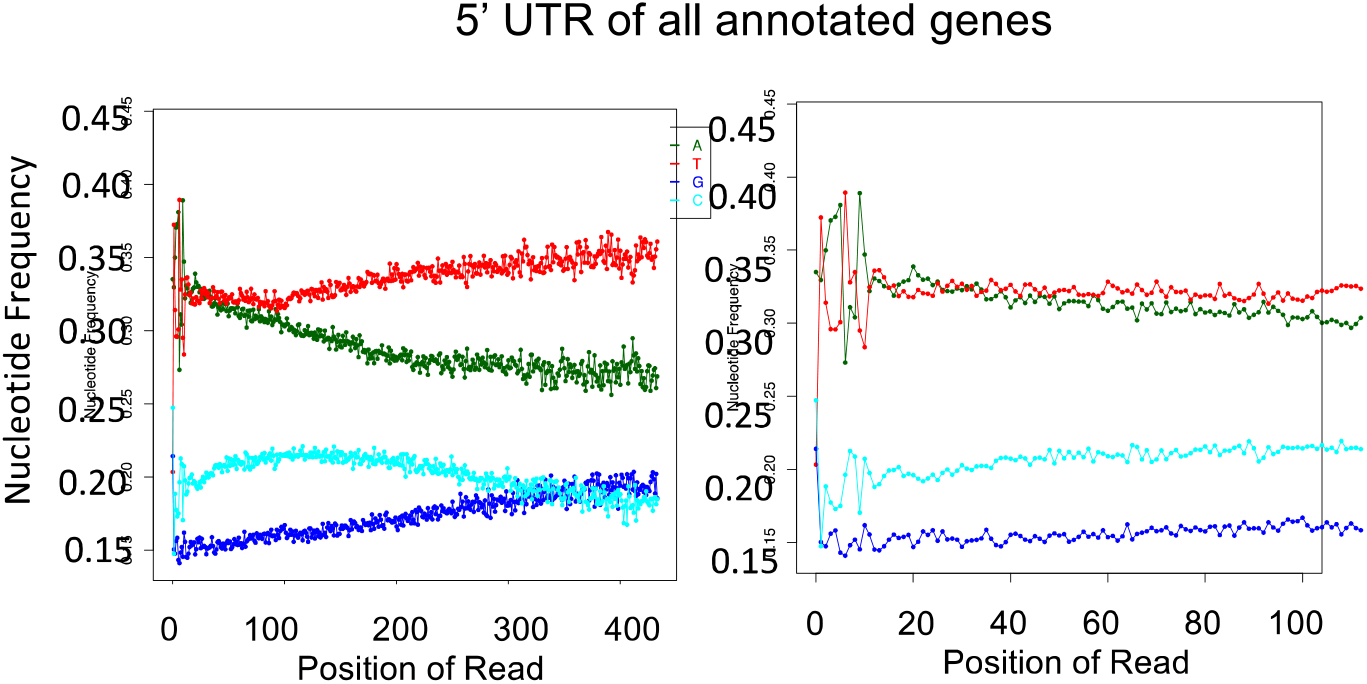
5’ UTR sequences of all annotated genes. The longest 5’ UTR for each gene were used to calculate the nucleotide frequency. The left panel is the distribution of nucleotides in the first 400 bp of the UTR region, and the right panel is the distribution of nucleotides in the first 100 bp of the UTR region.

